# Mesenchymal stem cells with overexpression of Angiotensin-converting enzyme-2 improved the microenvironment and cardiac function in a rat model of myocardial infarction

**DOI:** 10.1101/2020.05.03.075283

**Authors:** Chao Liu, Yue Fan, Hong-Yi Zhu, Lu zhou, Yu Wang, Qing-Ping Li

**Author notes:** Corresponding author. (Q.P.Li), (C.L.) Address: Department of Pharmacology, Nanjing Medical University, 101 Longmian Avenue, Nanjing 211166, P.R.China.

## Abstract

**Background:** Angiotensin-converting enzyme-2 (ACE2) overexpression improves left ventricular remodeling and function in diabetic cardiomyopathy; however, the effect of ACE2-overexpressed mesenchymal stem cells (MSCs) on myocardial infarction (MI) remains unexplored. This study aimed to investigate the effect of ACE2-overexpression on the function of MSCs and the therapeutic efficacy of MSCs for MI.

**Methods:** MSCs were transfected with Ace2 gene using lentivirus, and then transplanted into the border zone of ischemic heart. The renin-angiotensin system (RAS) expression, nitric oxide synthase (NOS) expression, paracrine factors, anti-hypoxia ability, tube formation of MSCs, and heart function were determined.

**Results:** MSCs expressed little ACE2. ACE2-overexpression decreased the expression of AT1 and VEGF apparently, up-regulated the paracrine of HGF, and increased the synthesis of Angiotensin 1-7 in vitro. ACE2-overexpressed MSCs showed a cytoprotective effect on cardiomyocyte, and an interesting tube formation ability, decreased the heart fibrosis and infarct size, and improved the heart function.

**Conclusion:** Therapies employing MSCs with ACE2 overexpression may represent an effective treatment for improving the myocardium microenvironment and the cardiac function after MI.

## 1. Introduction

Cardiovascular disease remains a major cause of mortality in Western societies. [1]. Currently, the major therapies for myocardial infarction (MI) include drugs, percutaneous coronary intervention with stent implantation, trials of stem cell transplantation. Giving the great potential of self-renew and differentiation of stem cells, stem cell transplantation was expected to improve cardiac function via tissue regeneration [2,3]. Mesenchymal stem cells (MSCs) are widely used in clinical trials for MI treatment; whereas it is reported that MSCs give therapeutic benefits via paracrine of cytokines [4] more than differentiation into mature cardiomyocytes [5]. Thus, many efforts including pharmacological pretreatment [6,7] and genetic modification [8,9] have been made to enhance the paracrine effects of MSCs. For instance, we have reported that pretreatment of MSCs with angiotensin II (Ang II) enhanced paracrine effects, angiogenesis, gap junction formation and therapeutic efficacy for MI [7].

Angiotensin-converting enzyme-2 (ACE2), a key enzyme of renin-angiotensin system (RAS), converts Ang II to Angiotensin-(1-7) (Ang-(1-7)). The receptor of Ang-(1-7) is mas receptor[10]. In the progression of heart failure, progressively elevated Ang II in myocardium promotes cardiac fibrosis and remodeling. On the contrary, intravenous infusion of Ang-(1-7) preserved cardiac function, coronary perfusion, and aortic endothelial function in a rat model for heart failure [11]. It was reported that ACE2 mRNA remained elevated in all areas of the MI rat heart, and ACE2 protein localized to macrophages, vascular endothelium, smooth muscle, lung type II pneumocytes, and myocytes [12,13]. To increase the cardiac ACE2 expression and activity [14,15], researchers employed adenovirus vector to deliver the Ace2 gene and found that it ameliorates left ventricular remodeling and dysfunction in a rat model of MI [14]. However, the use of adenovirus vector showed issues like failure to maintain long-term expression and immunogenicity [16]. Besides, it was reported that heart block, ventricular tachycardia, and sudden death appeared in ACE2 transgenic mice with downregulated connexins [17]. This revealed a risk of adverse reaction of the direct ACE2-overexpression in myocardium.

Lentivirus vector (LVV) has high transduction efficiency in MSCs [18]. We supposed that transplantation of ACE2-overexpressed MSCs (LV-ACE2-MSCs) might provide better therapeutic efficacy than naive MSCs for MI through improvement of cardiac microenvironment. In this study, we investigated the therapeutic effects of ACE2 over-expression in MSCs for MI, and the mechanisms underlying this potential therapeutic strategy.

## 2. Material and methods

### 2.1 Isolation and culture of MSCs

MSCs were isolated from male Sprague–Dawley (SD) rats (60 to 80 g, provided by Experimental Animal Center of Nanjing Medical University, Nanjing, China). Rats were sacrificed by cervical dislocation. Then MSCs were generated by flushing the rats’ femurs with sterile Dulbecco’s modified Eagle’s medium (DMEM, Thermo Scientific, USA) and were plated in culture flasks with DMEM containing 10% fetal bovine serum (FBS, Hyclone, USA) as previously described [6]. The MSCs were then incubated at 37 °C for 3 d before the first change of medium. The mesenchymal population was isolated on the basis of its ability to adhere to the culture plate. The cells were subcultured at 80 to 90% confluence. Three to five passage-cells were used for the experiments.

### 2.2 Transfection of cells with lentivirus encoding rat Ace2

The MSCs were transfected with Ubi-MCS-3FLAG-SV40-ACE2 (LV-Ace2) vectors and Ubi-MCS-3FLAG-SV40-EGFP (LV-gfp) control vectors (produced by GeneChem Inc. Shanghai, China) as previously described [19]. Briefly, primary MSCs were seeded in plates (Costar, Corning, NY, USA) in DMEM. Twenty-four hours after seeding, MSCs were infected with LV-Ace2 and LV-gfp. The cells were incubated with the virus for at least 12 h in minimal culture medium. Then, unbound virus was removed and replaced with fresh medium. The cells were incubated for another 72 h before treatment. After determination of the effect of infection, multiplicity of infection (MOI) = 20 was chosen for the other experiments with sufficient over-expression of ACE2/EGFP and minimum harm to the infected cells.

### 2.3 Real-Time Quantitative PCR (qPCR)

Total RNA was extracted using the TRIzol reagent (Invitrogen Life Technologies, Gaithersburg, MD) and stored at −80°C. The SYBR Green Master was used according to the manufacturer’s instructions. The primer sequences (sense/antisense) were as follows: ACE2, 5’-ACAATTGTTGGAACGCTGCC-3’/5’-CGCTTCATCTCCCACCACTT-3’; Mas, 5’-ACGGAAGCATCACCAAGGAG-3’/5’-TCGAAAGCCACCCACATTCA-3’; β-actin, 5’-GCACCGCAAATGCTTCTA-3’/ 5’-GGTCTTTACGGATGTCAACG-3’. The specificity of the amplification product was determined by performing a melting curve analysis. Standard curves were generated for the expression of each gene by using serial dilutions of known quantities of the corresponding cDNA gene template. Relative quantification of the signals was performed by normalizing the signals of different genes with the β-actin signal.

### 2.4 Western blot analysis

Whole-cell extracts were prepared using lysis buffer (Beyotime, China). Equivalent amounts of protein were applied to 10% SDS-PAGE gels and transferred to a polyvinylidene difluoride (PVDF) membrane. The membrane was blocked in 5% BSA/TBST and incubated overnight at 4 °C with the following primary antibodies: nitric oxide synthase 1 (NOS1), NOS3, STAT3, VEGF, ACE, AT1, GAPDH (Santa Cruz Biotechnology), and ACE2 (Abcam). The membranes were exposed to HRP conjugated secondary antibodies for 2 h at room temperature, and protein expression was detected using an enhanced chemiluminescence (ECL) western blot detection reagent (Pierce).

### 2.5 Assay of ACE2 activity

The assays of ACE2 activity in cells or tissues are conducted according to the instructions of SensoLyte^®^ 390 ACE2 kit.

### 2.6 Proteome profiler array assay

The Proteome Profiler Rat Adipokine Array Kit (ARY016, R&D Systems) was used to analyze molecular changes in cell culture media according to the manufacturer’s instructions. Visualization and quantitation of the detected spots were performed using the Image J software. Two different batches of MSCs were tested, with each totaling four replicate dotblots. Dotblot pixel density values were corrected by normalizing each dotblot to the highest average control and subtracting the average pixel density values of negative control dotblots in each blot.

### 2.7 Radioimmunoassay and Enzyme-linked immunosorbent assay (ELISA)

The supernatants of pretreated MSCs were collected at different time points and that of untreated MSCs were also collected. Ang II concentration was measured using radioimmunoassay kit (D02PJB, Beijing north institute of biological technology, Beijing, China). Ang-(1-7) concentration was measured using ELISA kit (S-1330, Bachem, CA, USA)

### 2.8 Co-culturing and mixed-culturing of MSCs and H9c2 cells MSCs

MSCs and H9c2 cells were co-cultured using a Transwell^®^ 3450 cell culture cluster (Corning, NY, USA). The MSCs and H9c2 cells were seeded in the well or in the chamber respectively, according to different experiments. Cells were cultured in DMEM with 10% FBS. Ang II was added into the co-culturing system.

### 2.9 Cell morphological analysis

H9c2 cells were co-cultured with LV-GFP-MSCs and LV-ACE2-MSCs, respectively. Ang II (5 nM) was added into the co-culture system at the beginning, then after 12 hrs, the system was exposed to hypoxia for 10 hrs. H9c2 cells were stained using Hoechst 33342 staining kit and photographed under microscopy (Axio Vert.A1, Zeiss). The Cell apoptosis analysis was done by cell apoptosis-Hoechst staining kit (Beyotime Institute of Biotechnology, China)

### 2.10 Tube formation assay

Matrigel-induced tube formation using and MSCs were performed to assess the angiogenic activity of MSCs. Briefly, Matrigel (growth factor reduced, BD Biosciences, USA) was diluted 1:1 with PBS and added to 96-well plates in a volume of 50 μL per well. Plates were held at 37 °C for 30 min into form a gel layer [7]. MSCs and H9c2 cells were seeded in the well and the transwell respectively in six-well plates, and treated using Ang II. MSCs (10,000 cells/well) were then seeded in matrigel-precoated 96-well plates. The cells were incubated at 37 °C and observed after 12 hrs and 36 hrs. The number of tubes was counted under microscopy (Axio Vert.A1, Zeiss). At least five wells were viewed, and the number of tubes/well was counted and averaged.

### 2.11 Rat MI model

The female SD rats weighing 220 to 250 g were anesthetized with chloral hydrate (300 mg/kg, Sinopharm Chemical Reagent, Shanghai, China) via intraperitoneal injection (IP). Experimental MI was induced by permanent ligation of the left anterior descending artery (LAD), as previously described [7]. The development of MI was checked using electrocardiogram (ECG) 30 min after ligation and confirmed with echocardiography 2 d after the surgery.

### 2.12 MSC transplantation

Rats were grouped (n = 5) after LAD ligation (except sham group) to receive 80 μL DMEM without cells (model group) or with LV-GFP-MSCs (LV-GFP-MSC group), or LV-ACE2-MSCs (LV-ACE2-MSC group). In groups of LV-GFP-MSC and LV-ACE2-MSC, each rat received 4 injections of MSCs (total of 1×10^6^ cells/heart) into the border zone of the infarcted area 1 h after the ligation.

### 2.13 Histological analysis of the infarcted rat heart

The rats were sacrificed by chloral hydrate (300 mg/kg IP) followed by cervical dislocation 30 d after cell transplantation. Heart sections embedded in paraffin were cut into 4-μm slices. The structure of the heart was shown on hematoxylin and eosin (H&E) stained slides. The size of the fibrosis area was assessed using Masson’s trichrome staining. Measurements were performed on 3 separate sections of each heart, and the averages were used for statistical analysis. Additionally, the size of the infarct area was determined using triphenyl tetrazolium chloride (TTC) staining. The heart was sectioned in 2-mm slices and stained with 2% TTC for 20 minutes. For picrosirius red staining, sections were treated with 1% acetic acid for 2 min, stained by 0.1% Sirius Red for 20 min, then serially dehydrated in ethanol and xylene, and mounted. Images of the slices were captured and quantified using image analysis software (Image Pro Plus 6).

### 2.14 Immunohistochemical analysis

The rat hearts were cryopreserved in OCT compound (Tissue-Teck, Sakura, Torrance, California) and sliced transversely from the apex to the basal part of the left ventricle using a cryostat with an 8-μm thickness, primary antibodies against ACE2 (Abcam) were used and were detected with anti-rabbit IgG HRP (Abcam). Four tissue sections of each heart (5 hearts in each group) were randomly selected; Images were recorded with microscopy (Axio Vert.A1, Zeiss) and analyzed using Image Pro Plus 6.

### 2.15 Functional assessment of the infarcted rat heart with echocardiography

Transthoracic echocardiography was performed at 2 and 30 d after coronary ligation, as described previously [7], with an echocardiographic system (the Vevo^®^ 2100 system, VisualSonics, Canada) equipped with a 20-MHz linear-array transducer. Left ventricular (LV) end-diastolic and end-systolic dimensions (LVED,d and LVED,s) were measured according to the cutting-edge method of the American Society of Echocardiography. The LV percent fractional shortening (FS) and percent ejection fraction (EF) were automatically calculated by the Vevo^^®^^ 2100 system. The ΔEF and ΔFS values were calculated by comparing the data at 30 and 2 d.

### 2.16 Statistical analysis

All data are presented as mean ± SEM. Student’s *t*-test was used for two-group comparisons and ANOVA followed by an unpaired Student’s *t*-test was used for multiple group comparisons. A value of P < 0.05 was considered significant, and all of the data were analyzed using GraphPad Prism 5 software (GraphPad Prism Institute Inc).

## 3. Results

### 3.1 MSCs express little ACE2, which does not change by Ang II treatment

Firstly, we assessed the mRNA expression of Ace2 and Mas genes in MSCs using real-time qPCR. The results showed that the MSCs expressed little Ace2 (Ct = 33.06 ± 0.29, n=4) and a low level of Mas (Ct = 31.22 ± 0.15, n=3) gene (Figure S1A). Secondly, we tested whether Ang II pretreatment could up-regulate the expression of ACE2. MSCs were treated using 100 nM Ang II for 0 h (control), 6 h, 12 h, 24 h. Western blot analysis showed that MSCs expressed little ACE2, and Ang II pretreatment did not up-regulate ACE2 expression (Figure S1B).

### 3.2 ACE2 over-expression and MSCs proliferation

Lentivirus-mediated transduction and expression of ACE2 were confirmed with real-time qPCR and Western blot. PCR data indicated that expression of Ace2 was 7000-fold higher in LV-ACE2-MSCs (Figure S2A). LV-ACE2-MSCs also exhibited higher levels of ACE2 protein (Figure S2B). Moreover, the activity of over-expressed ACE2 in MSCs was determined using ACE2 activity kit. LV-ACE2-MSCs showed an about 40-fold higher level of activity (Figure S2C). To evaluate the impact of lentivirus infection or Ace2 over-expression on MSCs proliferation, cell viability was determined using CCK-8 kit. LV-GFP-MSCs showed a higher proliferation than naive MSCs, whereas Ace2 over-expression attenuated it (Figure S2D). When 5 nM Ang II, as a substrate, was supplied to MSCs for 24 h, LV-ACE2-MSCs showed a lower proliferation than LV-ACE2-MSCs (Figure S2E).

### 3.3 The impact of ACE2 over-expression on the expression of RAS components, nitric oxide synthase and paracrine factors

We determined the expressions of RAS components, nitric oxide synthase (NOS) and related key molecules at 3 d after MSCs were transfected with LV-gfp and LV-ace2. The expression of ACE2 in the LV-ACE2-MSC group was much higher than that in the LV-GFP-MSC group, whereas ACE did not change significantly. The expression of AT1 decreased apparently. The expression of nos3 was higher in the LV-ACE2-MSC group, whereas nos1 did not changed significantly. The expression of VEGF down-regulated apparently (Figure 1 A). To determine the paracrine factors, the rat adipokine array kit was used. The results showed that compared with LV-GFP-MSC group, the paracrine of VEGF, FGF-21, and IGF-I decreased in the LV-ACE2-MSC group, whereas the HGF and IGF-II increased. If 30% alteration was set as significance according to the instruction of the kit, HGF increased significantly (Figure 1 B).

**Fig. 1.**
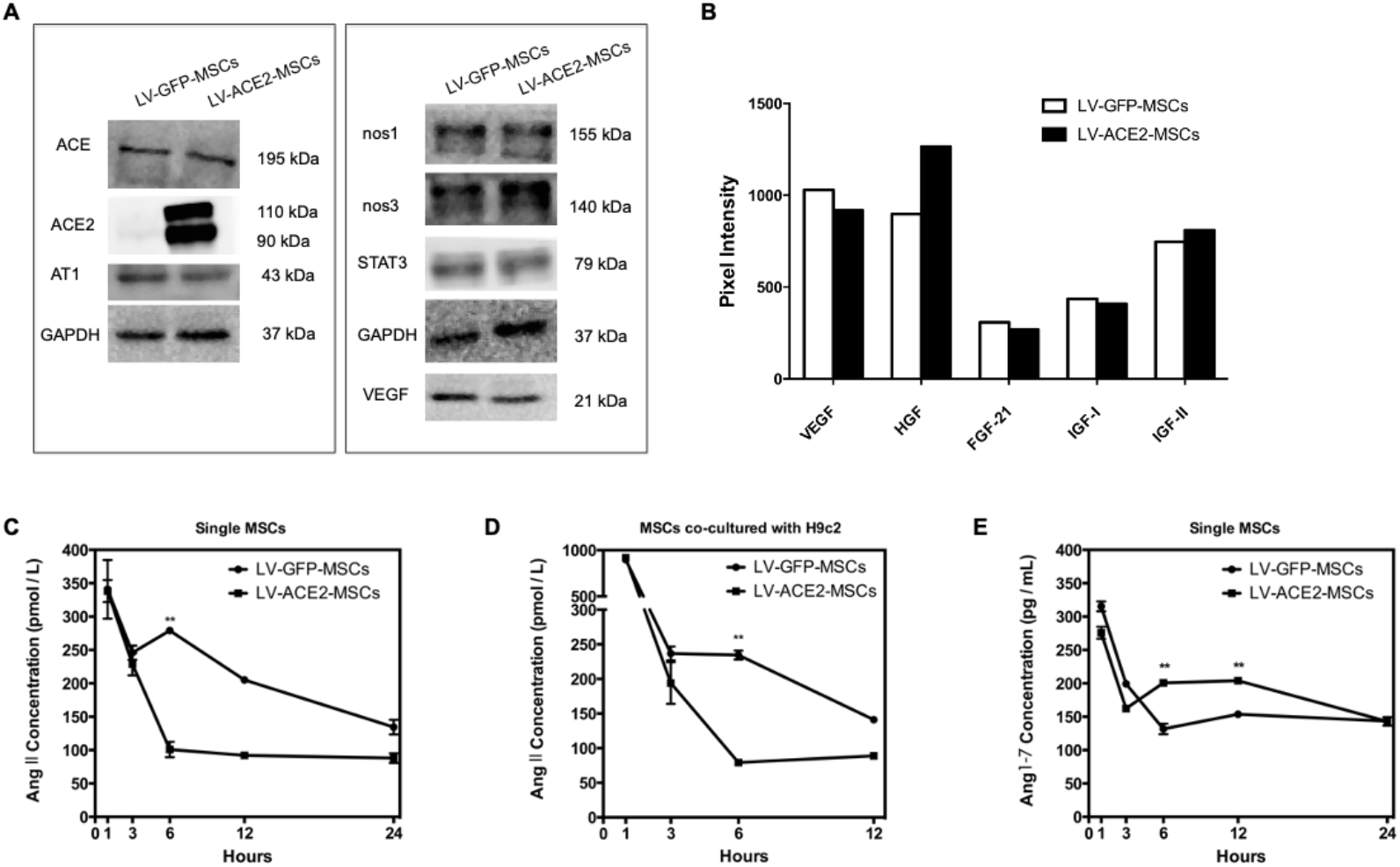
The influence of ACE2-overexpression on the RAS, NOS, and paracrine factors. (A) After MSCs were transfected with LV-gfp and LV-ace2, they were harvested and tested using western blotting (n= 5 in each group). (B) LV-GFP-MSCs and LV-ACE2-MSCs were co-cultured with H9c2 cells in DMEM (containing 5 nM Ang II) for 24 hrs. The culture media were then tested using Rat Adipokine Array Kit. The average signal (pixel density) of the pair of duplicate spots representing each protein was determined. (C) The concentrations of Ang II in single-cultured MSCs were determined by RIA. (D) The Ang II concentrations in MSCs co-cultured with H9c2 were determined. (E) The production of Ang-(1-7) was determined by EIA. The Ang-(1-7) concentration in the LV-ACE2-MSC group was significantly higher than that in the LV-GFP-MSC group at 6 h and 12 h. All values were expressed as mean ± SEM, **P < 0.01, n=3.

### 3.4 The effects of ACE2 over-expression on the level of Ang II/Ang-(1-7)

The concentrations of Ang II and Ang-(1-7) in the MSC supernatants were determined using the radioimmunoassay (RIA) and enzyme immunoassay (EIA). The MSCs were single cultured or co-cultured with the H9c2 cells. The substrate, a concentration of 5 nM Ang II, was added into MSC supernatants, and concentrations were determined within 24 hrs (Figure 1C and D). The concentration of Ang II decreased to less than 1 nM within 1 h. At the time point of 6 h, the concentration of Ang II in the LV-ACE2-MSC group was significantly higher than that in the LV-GFP-MSC group, which suggested a reduction of Ang II degradation or even a short increasing of Ang II (Figure 1C). After Ang II was added into supernatants once, the concentration of Ang-(1-7) was determined within 24 hrs. The results showed that the concentration of Ang-(1-7) in the LV-ACE2-MSC group was significantly higher than that in the LV-GFP-MSC group at 6 and 12 h, and then they reached the same at around 24 h (Figure 1E).

### 3.5 The impact of ACE2 over-expression on cytoprotective effect and angiogenesis

To reveal the cytoprotective effect of LV-ACE2-MSCs on H9c2 cells under hypoxia, we assessed cell apoptosis and cell death of H9c2 cells that were co-cultured with MSCs, after the cells were exposed to hypoxia (0.5% O_2_) for 10 hrs. In the LV-GFP-MSC group, H9c2 cells showed nuclear condensation and fragmentation, obvious morphological characteristics of apoptosis, while the number of which decreased in the LV-ACE2-MSC group (Figure 2 A and B). The number of the dead cells in the culture media in the LV-ACE2-MSC group was less than that in the LV-GFP-MSC group. (Figure 2 C). To estimate the influence of ACE2 over-expression on the angiogenic activity of MSCs, tube formation assay was performed and the results showed that the number of tube formation in the Matrigel in the LV-GFP-MSC group was more than that in the LV-ACE2-MSC group at 12 h; however, on the contrary, the number of tube formation in the LV-ACE2-MSC group was significantly higher at 36 h (Figure 2 D and E).

**Fig. 2.**
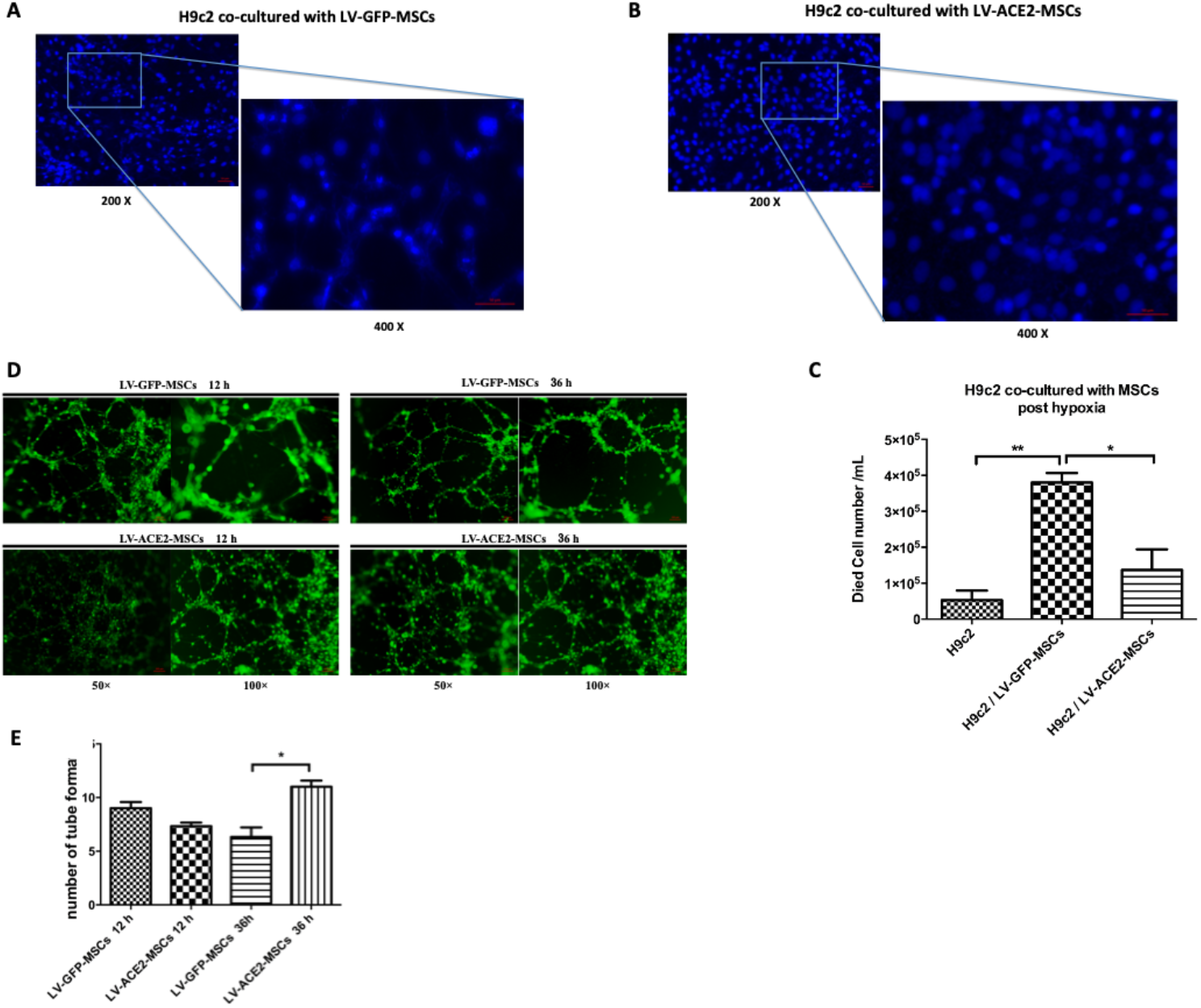
The impact of ACE2 over-expression on cytoprotective effect and angiogenesis. Examination of H9c2 cell apoptosis and death. (A and B) Morphologic images of H9c2 cell apoptosis. H9c2 cells were co-cultured with LV-GFP-MSCs and LV-ACE2-MSCs, respectively. Ang II (5 nM) was added into the co-culture system at the beginning, and after 12 hrs, the system was exposed to hypoxia for 10 hrs. The H9c2 cells were stained using Hoechst 33342 staining kit. (C) The H9c2 cell death was counted. The H9c2 cells without exposure to hypoxia were set as control. All values were expressed as mean ± SEM, **P < 0.01, *P < 0.05, n=3. Tube formation assay in vitro. (D) Representative pictures of tube formation of MSCs. Cells were treated with 5 nM Ang II for 12 hrs, and then cultured in the Matrigel (growth factor reduced) for 12 hrs and 36 hrs. (E) Statistical analysis of the number of tube formations in MSCs. At least five wells were viewed, and the experiments were repeated three times. All values were expressed as mean ± SEM, *P < 0.05, (n = 5 in each group).

### 3.6 LV-ACE2-MSCs transplantation increased heart ACE2 activity, decreased heart fibrosis and infarct size

Immunohistochemical results showed the expression of ACE2 in the model group, LV-GFP-MSC and LV-ACE2-MSC groups (Figure 3A). Fluorimetric analysis of ACE2 activity revealed that the level of ACE2 in the model group was higher than that in the sham group and was not significantly different from that in the LV-GFP-MSC group. The level of ACE2 in the LV-ACE2-MSC group was significantly higher than that in the LV-GFP-MSC group (Figure 3B). Picrosirius red–staining images showed that the collagen content (red) in the myocardium of LV-GFP-MSC group was lower than that in the model group, and the collagen content in the LV-ACE2-MSC group was the lowest (Figure 3C). TTC staining data showed that compared with the model group, transplantation of LV-GFP-MSCs decreased the infarct area, and LV-ACE2-MSCs transplantation showed even smaller infarct area (Figure 3 D and E).

**Fig. 3.**
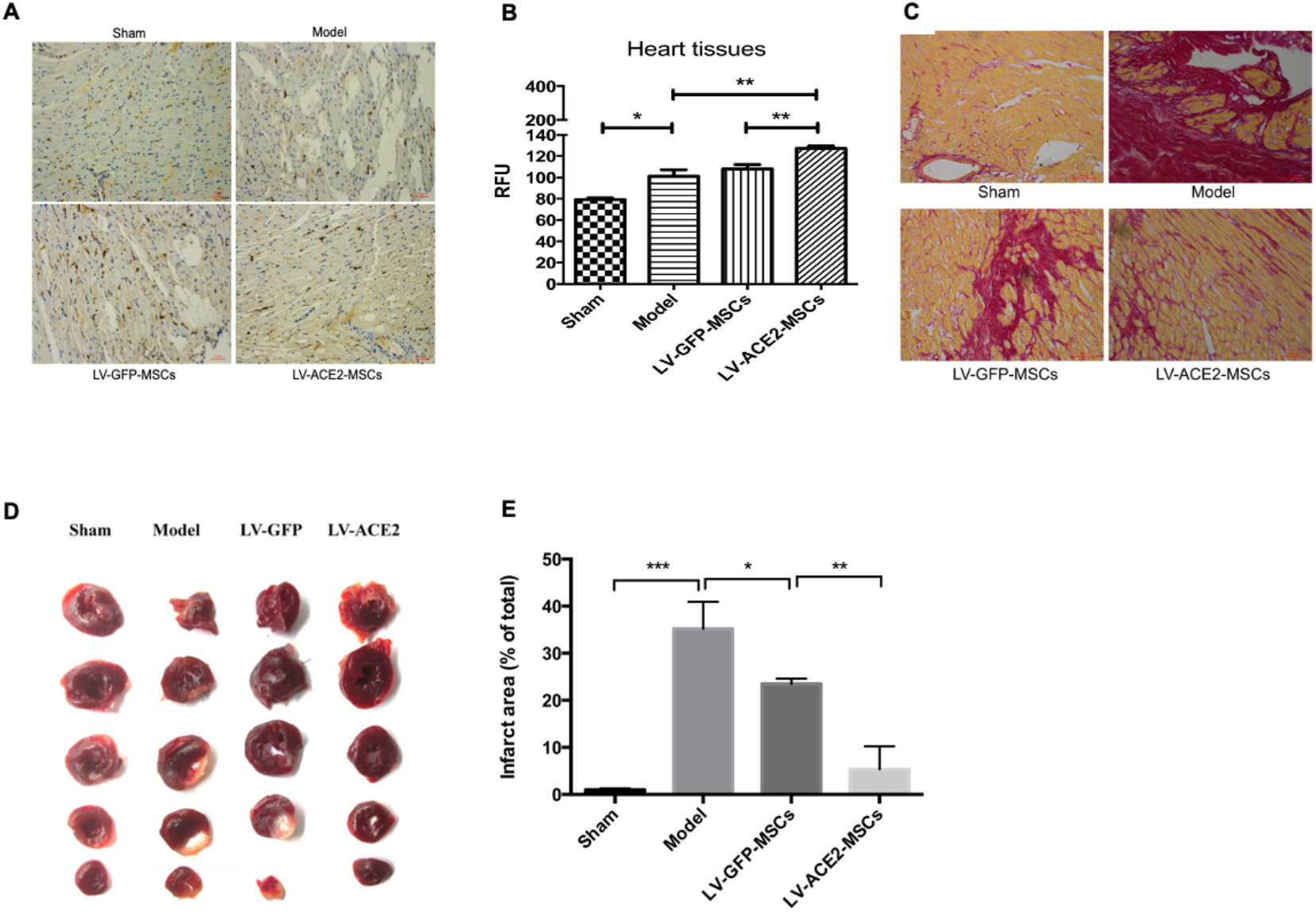
LV-ACE2-MSCs transplantation increased the heart ACE2 activity, decreased heart fibrosis and infarct size. (A) Representative images of ACE2 expression in ischemic heart tissues (×200), n=3. (B) Fluorimetric analysis of ACE2 activity in heart tissues using ACE2 Kit. All values were expressed as mean ± SEM, **P < 0.01, *P < 0.05, n=3. (C) Representative images of picrosirius red-staining of infarct heart. Cardiac muscle stained yellow, collagen stained red. The areas of collagen were observed (×200) and analyzed, n=3. (D) Representative TTC staining images. (E) Quantitative analysis of the heart infarct size in TTC staining images. All values were expressed as mean ± SEM, ***P < 0.01, **P < 0.01, *P < 0.05 (n=3 in each group).

### 3.7 LV-ACE2-MSCs exhibited better therapeutic effects in an MI model

To investigate the therapeutic effects of ACE2-overexpressed MSCs on ischemic heart, we transplanted ACE2-overexpressed MSCs into MI-model rats. Briefly, two days after LAD ligation and cell transplantation, rats in all groups except sham presented ischemic symptoms of persistent ST-segment elevation on ECG and EF<55% on echocardiography. There was no significant difference in the EF and FS percentages (%EF and %FS) among the groups that underwent LAD ligation. Thirty days after cell transplantation, echocardiography data showed that the LV anterior wall motion was obviously better in the LV-ACE2-MSC group than that in the LV-GFP-MSC group or the model group (Figure 4A). %EF and %FS were significantly higher in the LV-ACE2-MSC group (Figure 4B and C). Besides, the values of ΔEF and ΔFS in the LV-ACE2-MSC group versus LV-GFP-MSC group showed that transplantation of ACE2-overexpressed MSCs yielded better outcomes (Figure 4D and E).

**Fig. 4.**
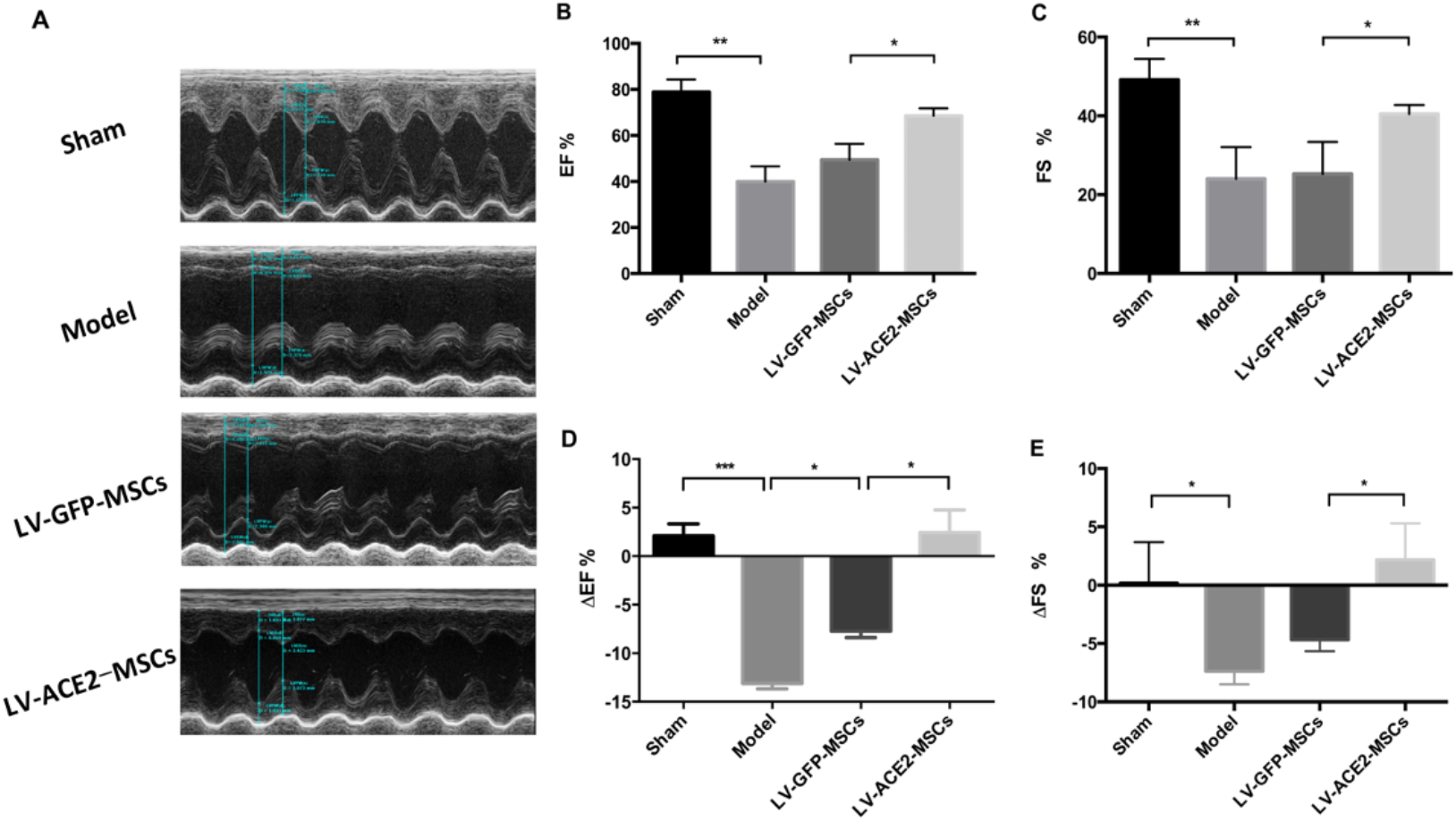
Evaluation of left ventricular function after cell transplantation. (A) Representative images of echocardiographic data that showed anterior wall motion (B and C) Echocardiography was performed at 30 d, %EF and %FS were automatically calculated. (D and E) The ΔEF and ΔFS values were calculated by comparing the data at 30 d and 2 d. All values were expressed as mean ± SEM, ***P < 0.001, **P < 0.01, *P < 0.05 (n=5 in each group).

## 4. Discussion

In this paper, we found MSCs expressed little ACE2 and the pattern of how ACE2 overexpression regulated the levels of RAS components and NOS. ACE2 over-expressed MSCs had a cytoprotective effect on cardiomyocyte in hypoxia and promoted angiogenesis probably through the up-regulation of HGF. This study was the first to reveal that ACE2 overexpression in MSCs improved left ventricular function in MI through increasing heart ACE2 activity, decreasing heart fibrosis, and infarct size. This study provided insight into the mechanism of the influence of ACE2-overexpression on the function of MSCs and on the therapeutic efficacy of MSCs for MI.

### 4.1 ACE2 over-expression and RAS components, NOS and paracrine factors

It has been reported that the RAS is active in the ischemic myocardium, especially the ACE activity and Ang II level is increasing [20,21]. Ang II is related with the acceleration of pathological process of MI. This study showed that ACE2 overexpression did not change the level of ACE, whereas down-regulated AT1 significantly. After the addition of a modest Ang II (5 nM), we found that Ang-(1-7) level was significantly elevated in the LV-ACE2-MSC group. Besides, we found that the over-expression of ACE2 in MSCs increased the expression of nos3, which was probably through Ang-(1-7), because Ang-(1-7) was reported to induce the up-regulation of nos3 via mas receptor [22]. Moreover, we found that ACE2 overexpression interestingly increased the HGF level in the cell co-culture medium instead of the VEGF, FGF-21, IGF-I, IGF-II. We have reported that Ang II treatment increased paracrine of VEGF, and this effect was probably attenuated by Ang-(1-7). We speculated that the increasing of HGF level was due to the paracrine of MSCs in the co-culture system, while the mechanism of which warrants further investigation. In a summary, we suggested that delivering of LV-ACE2-MSCs could improve the microenvironment of the ischemic myocardium.

### 4.2 The Ang II/ACE2/Ang-(1-7) levels and the cytoprotective effect

We found that ACE2 over-expression modestly decreased MSC proliferation (Fig. S2E) in the normal culture condition, and whether this affection was induced by ACE2 related pathway or by Ang-(1-7) remained to be elucidated. The dynamic of Ang II /Ang-(1-7) levels in the cell culture system was determined in this study, and according to the results (Fig. 1), we speculated that exogenous Ang II was degraded rapidly by neprilysin that is expressed in MSCs [23], at the same time, Ang II was converted into Ang-(1-7). In this process, the positive feedback phenomenon (Ang II level at 6 h time point) of Ang II [23] existed, which was abolished by ACE2 overexpression in the LV-ACE2-MSC group. The Ang-(1-7) level was significantly elevated for a period of at least 6 hrs in the LV-ACE2-MSC group. We found that compared with LV-GFP-MSCs, LV-ACE2-MSCs showed higher ability to protect cardiomyocytes from the hypoxia and low nutrition condition in vitro. We supposed that this effect is at least partially related to Ang-(1-7). As is mentioned, ACE2 overexpression interestingly increased the HGF level in the cell co-culture, it is probable that HGF also contributed to the cardiomyocytes protective effect of LV-ACE2-MSCs.

### 4.3 ACE2 over-expression and Angiogenesis

We have reported that Ang II pretreatment increased the angiogenesis activity of MSCs [7]. In this study, interestingly, we found that it was at 36 h, instead of 12h, that LV-ACE2-MSC group showed more tube formation than LV-GFP-MSC group, although the total amount of tubes at 36 h was less than that at 12 h. We suggested that these phenomena were the balance of Ang II and Ang-(1-7), and the difference of tube formation at 36 h might be related to the increasing of HGF. The HGF was reported to be an angiogenic growth factor, which accumulated during the cell culture [24] and induced angiogenesis without vascular inflammation, different from bFGF and VEGF [25].

### 4.4 LV-ACE2-MSCs transplantation and the outcomes

In the previous study, we have reported that pretreatment of MSCs with Ang II enhances paracrine effects, angiogenesis, gap junction formation and therapeutic efficacy for MI [7]. Although Ang II pretreatment raise the survival of MSCs, the effect of paracrine is short and limited [23]. The therapeutic effect of MSCs on MI may not last long. Thus, here we explored a potential long-term stem cell strategy for MI. The MSCs were a kind of good vectors to deliver the Ace2 gene into the ischemic myocardium, and Ace2 gene could express stably in the MSCs if the transplanted MSCs survived. In the ischemic myocardium, Ang II accelerated cardiomyocyte apoptosis and myocardium fibrosis could probably be attenuated by LV-ACE2-MSCs.

## 5 Conclusion

ACE2 over-expression in MSCs showed better therapeutic effects for MI through improving the myocardium microenvironment and the cardiac function.

## Declarations

### Ethics statement

This study was approved by the Committee on the Ethics of Animal Experiments of Nanjing Medical University and was in accordance with the *Guide for the Care and Use of Laboratory Animals* issued by the National Institutes of Health (NIH). Efforts were made to minimize the suffering of animals.

### Availability of data and materials

The data that support the findings of this study are available from the corresponding author upon reasonable request

### Competing interests

None declared

### Funding

This study was supported by the National Science Foundation of China [No.30973534 to Q-P.L., 81173052 to Q-P.L.]; and the Jiangsu College Graduate Research and Innovation Program [No.CXZZ13_0604 to C.L.].

### Authors’ contributions

C.L. and Q.P.L contributed the study design; C.L. and Q.P.L contributed the Fundings; C.L., Y.F., H.Y.Z, and L.Z. contributed the experiments. Y.W. contributed the paper revision.

## Acknowledgements

We thank Professor Dongya Zhu, leader of Jiangsu Genetic Engineering Drug Technology Center, for generously providing the gfp transgenic rats.

## Supplementary Information

**Fig. S1.**
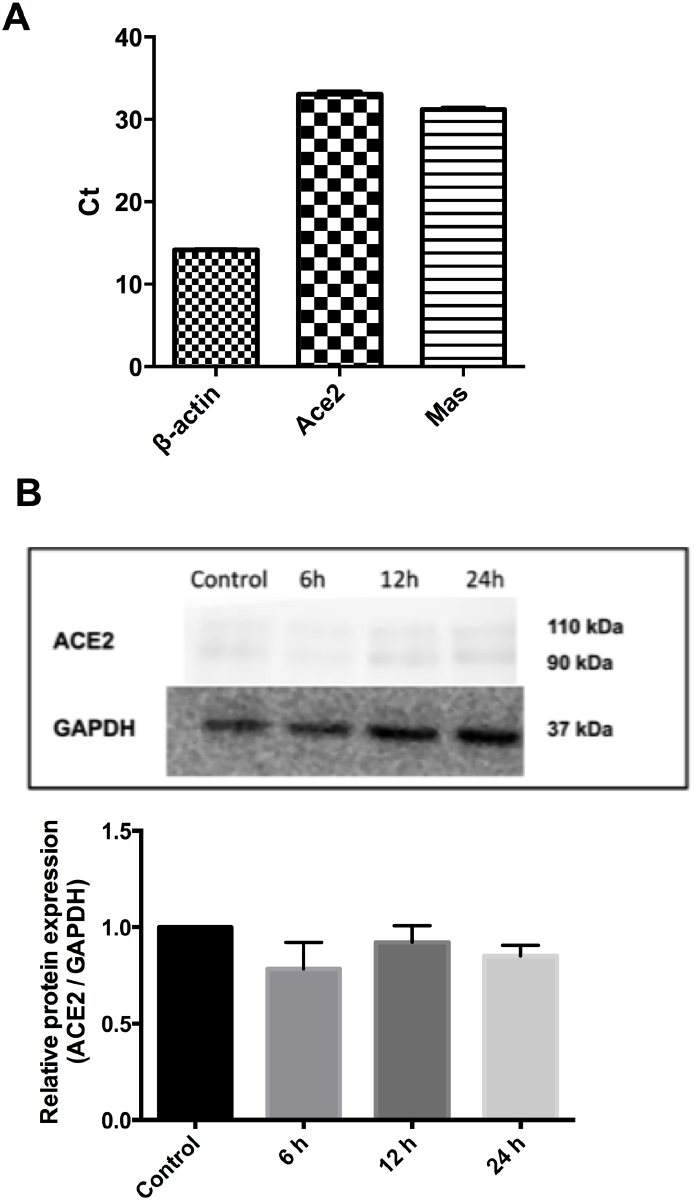
The MSCs expressed little ACE2, which did not change after Ang II treatment. (A) Examining mRNA expressions of the Ace2 and Mas genes in the MSCs by real-time qPCR. The MSCs were isolated from rats and the second passage was used for this experiment. n=3. (B) Analysis of the influence of Ang II treatment on the ACE2 expression. MSCs expressed little ACE2, and Ang II treatment did not up-regulate ACE2 expression. All values were expressed as mean ± SEM, P > 0.05. n=3.

**Fig. S2.**
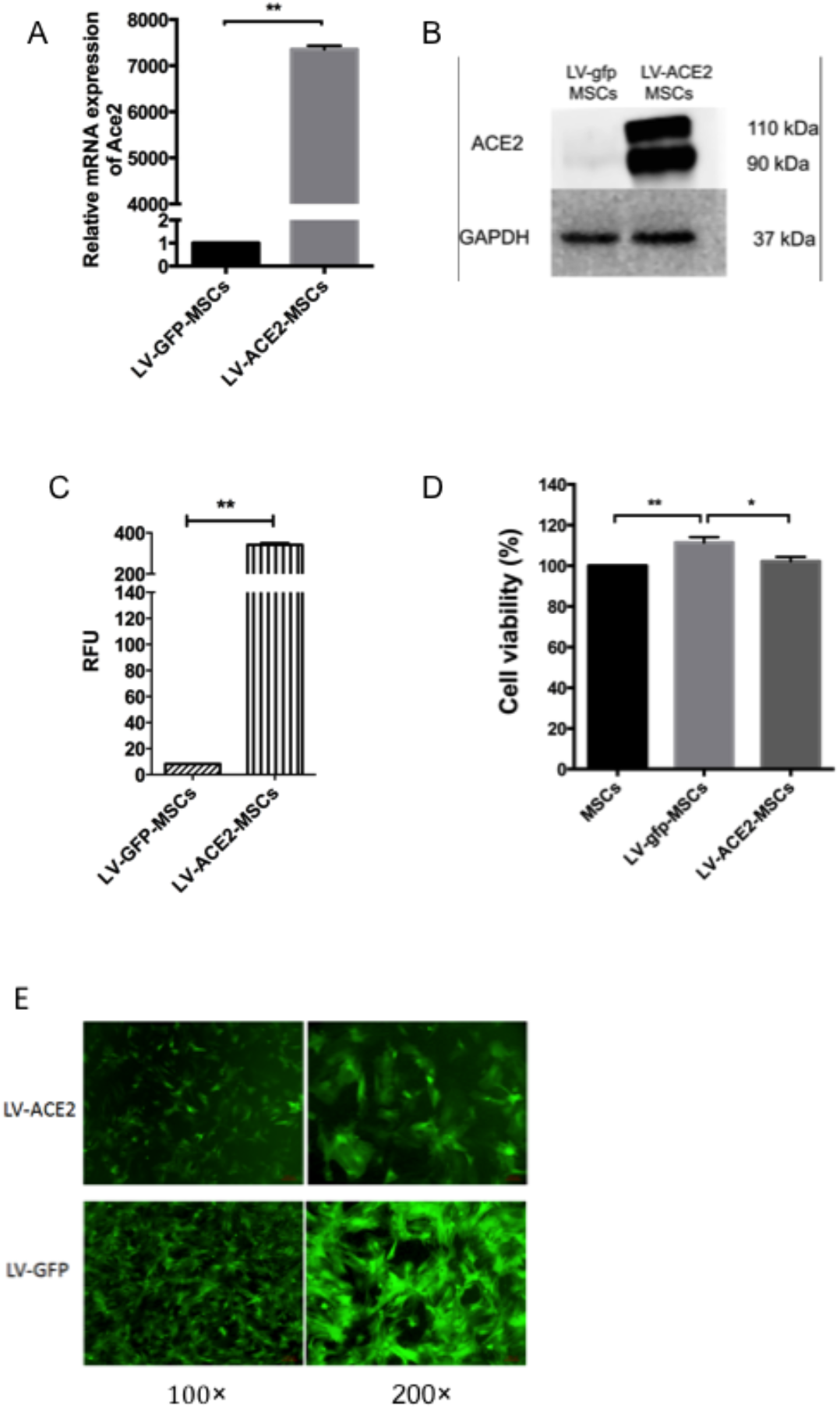
ACE2 over-expression and MSCs proliferation. The expression and activity of ACE2 in transfected MSCs were determined. (A) Real-time qPCR analysis of the Ace2 expression in MSCs. (B) Western blot analysis of the ACE2 expression in MSCs. (C) Fluorimetric analysis of the ACE2 activity using ACE2 Assay Kit. All values were expressed as mean ± SEM, **P < 0.01, n=3. (D) The cell viability was tested using CCK-8 kit. All values were expressed as mean ± SEM, **P < 0.01, *P < 0.05, n=5. (E) The observation of LV-MSCs proliferation after the addition of 5 nM Ang II substrate for 24 hrs. n=5.

## Notes

### Competing Interest Statement

The authors have declared no competing interest.

